# Chromosome-level reference genome assembly of the Saimaa ringed seal (*Pusa saimensis*) – an ancient glacial relict landlocked pinniped

**DOI:** 10.64898/2026.08.13.744633

**Authors:** Martin Grethlein, Zsófia Fekete, Steffi Goffart, Angelika Kiebler, Mervi Kunnasranta, Marja Niemi, Danilo F. Santoro, Gerrit Wehrenberg, Sven Winter, Stefan Prost, Jaakko Pohjoismäki

**Author notes:** These authors contributed equally (Stefan Prost, Jaakko Pohjoismäki).

## Abstract

We present a high-quality chromosome-level reference genome for the Saimaa ringed seal (*Pusa saimensis*), an endangered freshwater pinniped endemic to Lake Saimaa, Finland. The assembly spans 2.353 Gb and comprises 15 autosomes together with the X and Y sex chromosomes. Using Oxford Nanopore Technologies (ONT) long-read sequencing and Hi-C scaffolding, we achieved a telomere-to-telomere assembly for all chromosomes, except the Y chromosome. Genome annotation identified approximately 21,800 protein-coding genes, consistent with other mammalian genomes. Assembly completeness was high, with BUSCO analysis recovering 99.6% of expected complete single-copy genes (98.2% single-copy and 1.3% duplicated).

Comparative analyses revealed a highly conserved chromosomal architecture, with only minor syntenic differences relative to other pinniped chromosome-level assemblies. Previously described cytogenetic fusion events in Phocidae were confirmed (chromosomes 2 and 7). A translocation between chromosomes 6 and 7 distinguishes phocids from the otariids. In general, more distantly related taxa exhibit an increasing degree of intrachromosomal rearrangements. Notably, we identified a large intrachromosomal rearrangement on chromosome 2 that appears specific to the Saimaa ringed seal.

Phylogenomic analysis based on 9,226 single-copy orthologues placed the Saimaa ringed seal as a sister lineage to the Baltic ringed seal (*Pusa hispida botnica*), while confirming also other established evolutionary relationships among pinnipeds. Comparative gene family analysis between the Saimaa ringed seal and the closely related grey seal (*Halichoerus grypus*) revealed lineage-specific differences driven by a limited number of gene families. In the Saimaa ringed seal, expansions were observed in ion transport, cytoskeleton, and regulatory genes, potentially reflecting adaptation to freshwater conditions. In contrast, the grey seal showed expansions in olfaction, immune-and spermatogenesis-associated gene families, including MAGE/MIA genes, consistent with differences in ecology and mating systems.

This reference genome provides an important resource for studies of pinniped genome evolution, as well as conservation and population genomics of the Saimaa ringed seal, facilitating future work on genetic diversity, inbreeding, mutational load and adaptive potential in this highly endangered species.

## 1 Introduction

The Saimaa ringed seal, *Pusa saimensis* (Nordqvist, 1899), is a unique endemic freshwater pinniped confined to Lake Saimaa in southeastern Finland (Kunnasranta et al., 2021)(Figure 1). Until recently, it was considered to be a subspecies of the ringed seal, *Pusa hispida* (Schreber, 1775), the most widespread pinniped on the northern hemisphere with a circumpolar distribution (Berta & Churchill, 2012). It was thought that after initial dispersal into the Baltic basin of Fennoscandia at the end of the ice age, ringed seals had diverged into three subspecies in this area: the Baltic ringed seal, *P. h. botnica* (Gmelin, 1788), inhabiting the northern Baltic Sea, the Ladoga ringed seal, *P. h. ladogensis* (Gmelin, 1788), confined to Lake Ladoga, and the Saimaa ringed seal in Lake Saimaa (Ukkonen et al., 2014). However, recent genomic evidence indicates that the lineage leading to the ancestor of the Saimaa ringed seal diverged from other ringed seals at least 60,000 years ago (Löytynoja et al., 2025), which not only substantially predates the isolation of the Baltic and Ladoga populations but also the whole post-glacial history of the Baltic basin (Bjorck, 1995; Patton et al., 2017). It has been suggested that the Saimaa ringed seal represents the last surviving lineage of ringed seals that inhabited the extensive glacial lakes of northwestern Russia (Löytynoja et al., 2025), formed during the early phases of the Weichselian glaciation (Krinner et al., 2004; Mangerud, Astakhov, Jakobsson, & Svendsen, 2001; Mangerud et al., 2004), and subsequently dispersed into the Baltic basin following the retreat of the Fennoscandian ice sheet (Bjorck, 1995; Johansson, 2007; Patton et al., 2017). In recognition of its distinct evolutionary trajectory, including unique morphological adaptations to freshwater ecosystems (Hyvärinen & Nieminen, 1990; Löytynoja et al., 2025; Nihtilä & Laakkonen, 2025), the Saimaa ringed seal has been elevated from subspecies to full species (Löytynoja et al., 2025), a status that has also been formally accepted by the Society for Marine Mammalogy (Committee_on_Taxonomy, 2025). As such, the species designation provides a more solid framework for conservation, management, and comparative genomic research.

**Figure 1.**
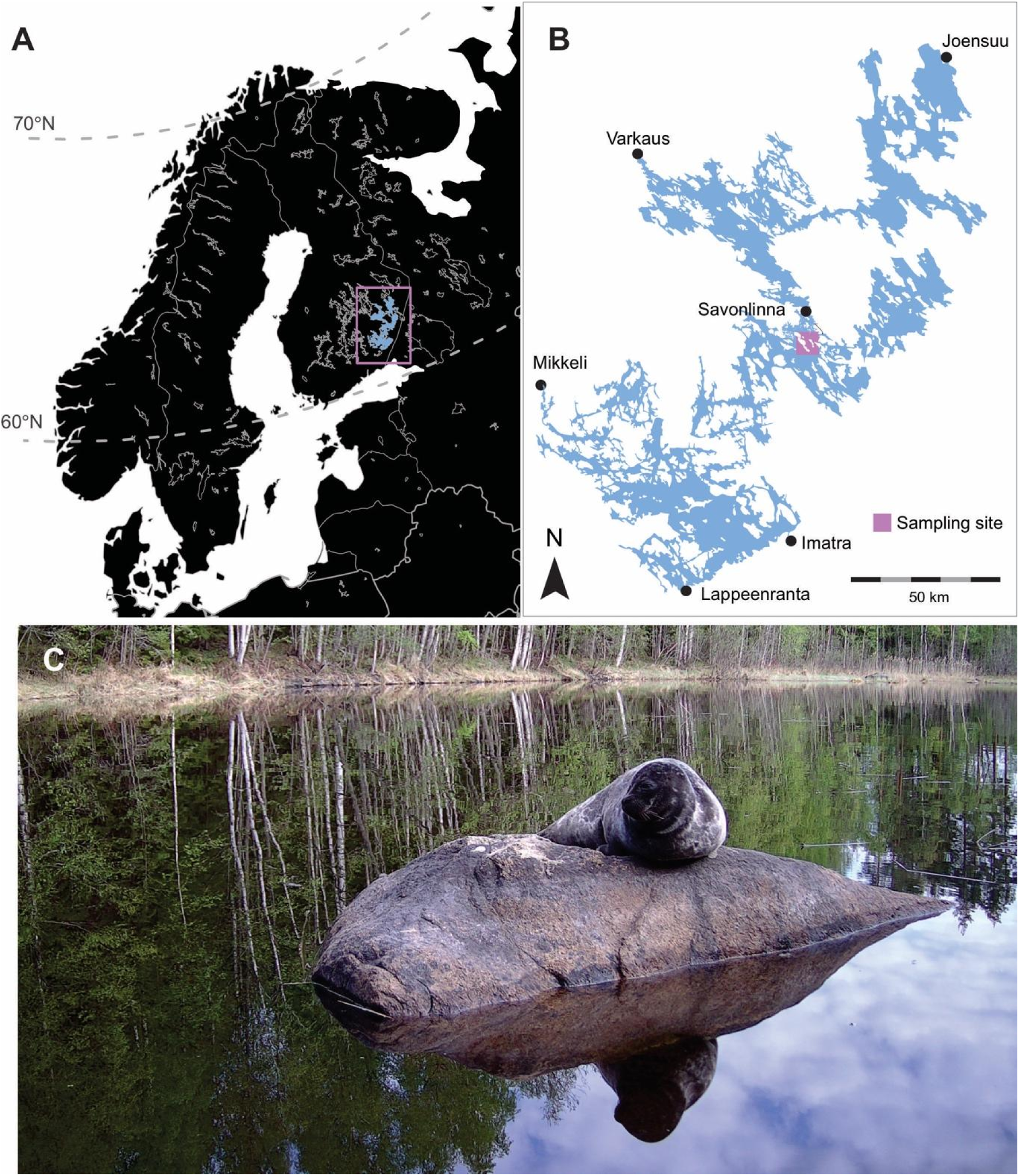
Saimaa ringed seal sampling for the reference genome. (A) The species inhabits Lake Saimaa, a large (ca. 4,400 km^2^), geographically complex freshwater system located in southeastern Finland (boxed area). (B) Although the species occurs throughout much of the lake, the majority of the 500+ individuals are concentrated in its central region. The largest subpopulation occurs in Pihlajavesi, south of the town of Savonlinna, where the reference genome individual also originates from. The sampling site and major towns are highlighted. (C) The sampled male (Phs449) hauled out on a rock. Photo: Saimaa ringed seal research group, University of Eastern Finland.

Today, the Saimaa ringed seal is among the most genetically depauperate mammalian species worldwide (Löytynoja et al., 2025; Nyman et al., 2014; Olkkonen & Löytynoja, 2023; Valtonen, Palo, Ruokonen, Kunnasranta, & Nyman, 2012), and it is classified as “Endangered” (EN) on the Red List of Threatened Species of Finland (Liukko et al., 2019; Sipilä, 2016). Long-term isolation, founder effects, repeated demographic bottlenecks, and a severe human-induced population collapse during the 19^th^ and 20^th^ centuries have all contributed to reduced genetic diversity (Heino et al., 2023; Kunnasranta et al., 2021; Löytynoja et al., 2025; Nyman et al., 2014; Olkkonen & Löytynoja, 2023; Olsen et al., 2025; Palo, Hyvärinen, Helle, Mäkinen, & Väinöla, 2003; Sundell et al., 2023; Valtonen et al., 2014; Valtonen et al., 2012). The population, which might have declined to fewer than 100 individuals in the 1960s has recovered to over 500 animals in 2025 (Metsähallitus, 2025), following legal protection and intensive conservation measures (Kunnasranta et al., 2021). However, the species remains under threat of extinction, and distribution is fragmented into several partially isolated subpopulations within the labyrinthine lake system, resulting in restricted gene flow, local differentiation, and elevated inbreeding pressure (Olkkonen & Löytynoja, 2023; Sundell et al., 2023). At the same time, this subdivision has preserved unique genetic variants within different lake basins (Löytynoja et al., 2025; Sundell et al., 2023; Valtonen et al., 2014), representing important evolutionary potential for future adaptations.

High-contiguity reference genomes provide essential resources for investigating genome structure, evolutionary history, and the functional basis of phenotypic traits (Blaxter et al., 2022). They enable accurate detection and localisation of genetic variation, including single-nucleotide polymorphisms and structural variants (Hu et al., 2025; Kim, Oh, Kim, & Kim, 2026), as well as runs of homozygosity (Taylor, Manseau, & Wilson, 2025), allowing the reconstruction of demographic history and population structure (Genereux et al., 2020; Sozzoni et al., 2025). Beyond population genomics, chromosome-level assemblies facilitate comprehensive gene annotation and comparative analyses across species (Gaertner, Tapanainen, et al., 2025; Gaertner, Terzioglu, et al., 2025), supporting studies of gene family evolution, regulatory elements, and signatures of natural selection (A. Rhie et al., 2021). They also provide the foundation for functional genomic approaches, including transcriptomics, epigenomics, and the identification of candidate genes underlying adaptation, disease susceptibility, reproduction, metabolism, and environmental resilience (Genereux et al., 2020). In conservation contexts, such reference genomes enable the development of precise genetic tools for monitoring diversity, relatedness, inbreeding, and adaptive potential, thereby supporting evidence-based management and long-term species persistence (Wehrenberg et al., 2025). Prior to the Saimaa ringed seal genome assembly presented here, chromosome-level genomes have been available for the Baikal seal (*Pusa sibirica*, GenBank reference GCA_028975605.1) (Yakupova et al., 2023), grey seal (*Halichoerus grypus*, GenBank reference GCF_964656455.1; no publication), Mediterranean monk seal (*Monachus monachus*, GCA_976986665.1; no publication), Weddell seal (*Leptonychotes weddellii*, GCA_054177405.2) (Noh et al., 2022), Hawaiian monk seal (*Neomonachus schauinslandi*, GCA_002201575.2)(Mohr et al., 2022) and the Californian sea lion (, GCA_009762305.2)(Peart et al., 2021). The previously available draft assembly for the Saimaa ringed seal (GCA_947044825.1) was generated at contig level. The unannotated assembly consisted of 2,183 contigs/scaffolds, with N50 of 3.7Mb and L50 of 183. While relatively contiguous for a draft assembly, the lack of chromosome-level scaffolding limited its utility for chromosome-scale genomics and structural analyses.

Here, we present a chromosome-level reference genome for the Saimaa ringed seal generated from a male individual. The assembly spans 2.353 Gb and comprises 15 autosomes and the X and Y sex chromosomes, containing approximately 21,800 annotated genes, as well as the 16,821 bp mitochondrial genome. The assembly forms a critical foundation for future studies on population genomic analyses of long-term isolation, small population size, adaptation to freshwater conditions, the identification of functional or adaptive variation and the development of genetic tools to support evidence-based conservation management of this evolutionarily exceptional freshwater seal.

## 2 Materials and Methods

### 2.1 Sampling, cell line isolation and DNA extraction

For this study, we used a cell line generated from a young male Saimaa ringed seal (“Tuukka”, individual code Phs449, Figure 1C). This individual, like all Saimaa ringed seals, can be uniquely identified by its permanent and distinctive fur patterns (Koivuniemi, Auttila, Niemi, Levänen, & Kunnasranta, 2016). The seal was live-captured in May 2023 in the Pihlajavesi basin of central Lake Saimaa (approx. WGS84 N61.8, E28.9) as part of a conservation translocation programme aimed at enhancing gene flow among subpopulations within the lake (Sundell et al., 2023). Due to the endangered status of the species, precise sampling coordinates are withheld and spatially generalized in accordance with conservation data sensitivity guidelines.

During the health assessment of the individual, a small skin biopsy was collected from the hind flipper under local anaesthesia by a licensed veterinarian. All animal handling and sampling procedures were conducted under permits ESAVI/34853/2022 and ESAELY/1035/2022 granted by the Finnish Regional State Administrative Agency (AVI; Animal Experiment Board, ELLA) and the South-Savo Centre for Economic Development, Transport and the Environment, respectively. Sampling adhered to the ARRIVE guidelines (Percie du Sert et al., 2020). The biopsy was used to establish a primary fibroblast cell culture following standard protocols for isolation and expansion of mammalian dermal fibroblasts (Reichel et al., 2026; Seluanov, Vaidya, & Gorbunova, 2010). The primary (non-transformed) cell line has been deposited as cryopreserved living cells under sample code ZFMK-TIS-133860 in the biobank of the Stiftung Leibniz-Institut zur Analyse des Biodiversitätswandels, Zoologisches Forschungsmuseum Alexander König (ZFMK), Bonn, Germany.

The cell line was immortalised using transformation with the plasmid pBSSVD2005 containing the SV40 large T-antigen (kind gift from David Ron; Addgene plasmid # 21826; http://n2t.net/addgene:21826; RRID: Addgene_21826). Cells used for DNA extraction represented passage number 10, counted from the isolation of primary fibroblasts from the biopsy. Cell lines below passage 20 are considered low passage and are expected to retain genomic integrity with minimal culture-induced alterations (Briske-Anderson, Finley, & Newman, 1997). High-molecular-weight genomic DNA was purified using the phenol:chloroform:isoamyl alcohol (IAA) extraction method (Sambrook & Russell, 2006). Briefly, fibroblast cells were cultured to confluency in a 10 cm tissue culture dish, detached using 10 mM EDTA in PBS, pelleted by centrifugation, and washed twice with phosphate-buffered saline (PBS). The cell pellet was resuspended in 2 ml of DNA extraction buffer (25 mM EDTA pH 8.0, 75 mM NaCl, 1% SDS, RNase A) and incubated with Proteinase K to ensure complete protein digestion. Following lysis, genomic DNA was extracted by sequential phenol:chloroform:IAA purification steps (Sambrook & Russell, 2006). DNA was ethanol-precipitated, washed, and resuspended in nuclease-free TE buffer.

### 2.2. Library preparation and sequencing

We prepared two sequencing libraries using Oxford Nanopore Technologies’ (ONT) Ultra-Long DNA Sequencing Kit (SQK-ULK114). As the DNA was extracted as detailed above, library preparation started at the tagmentation step. The protocol was modified as follows: During tagmentation, time on ice was increased to 15 minutes and incubation time for overnight elution after clean-up was set to 22 hours. The resulting libraries were sequenced on the PromethION P2 Solo (ONT, United Kingdom) platform, using R10.4.1 flow cells. The prepared libraries were split into three consecutive loads, each of which was sequenced for 21 hours, followed by a flush using ONT’s Flow Cell Wash Kit (EXP-WSH004). Incubation time with the Flow Cell Wash Mix was increased from one to three hours. Reloading the ultra-long DNA library after the flush was performed according to the manufacturer’s guidelines specified in the Ultra-Long DNA Sequencing Kit.

In addition, a proximity ligation library was generated from the same individual using the EpiTect-Hi-C Kit (Qiagen, Hilden, Germany, Cat. No. 59971) according to the manufacturer’s guidelines. The Hi-C library was sequenced by Novogene (Munich, Germany) on an Illumina NovaSeq X platform (Illumina Inc., San Diego, CA, USA).

### 2.3. Sequence data processing, assembly, scaffolding and curation

The raw ONT PromethION P2 Solo sequencing reads were basecalled using the super high accuracy basecalling model and duplex calling in Dorado 0.7.2 (https://github.com/nanoporetech/dorado). For assembly, we retained all reads with Phred quality scores >8 and included lower-quality reads when their lengths exceeded 5 kb, resulting in 60.6 Gb. Read quality metrics were assessed using NanoPlot (De Coster & Rademakers, 2023).

Assemblies of the nanopore data were carried out using Flye v2.9.3-b1797 (Kolmogorov, Yuan, Lin, & Pevzner, 2019) and Hifiasm v0.19.6-r597 (Cheng, Concepcion, Feng, Zhang, & Li, 2021), respectively. Assembly qualities were assessed using BUSCO v6.0.0 (Manni, Berkeley, Seppey, & Zdobnov, 2021). Continuity was assessed using gfastats v1.3.11 (Formenti et al., 2022).

Next, both assemblies were scaffolded using the Hi-C library (sequenced to 150 Gb). The scaffolding was performed using the Vertebrate Genome Project (VGP) Hi-C scaffolding pipeline (A. Rhie et al., 2021) and YaHS v1.2 (Zhou, McCarthy, & Durbin, 2023). In brief, the Hi-C data were first filtered using the filter_five_end.pl *perl* script (https://github.com/ignacio3437/HiC_mapping_pipeline/tree/master), followed by removal of PCR duplicates with *Sambamba* v1.0.1 (Tarasov, Vilella, Cuppen, Nijman, & Prins, 2015). Scaffolding was then carried out using YaHS v1.2 (Zhou et al., 2023). Gap closing was performed with *TGS-GapCloser* v2.0.0 (Xu et al., 2020).

Following a second round of quality assessment with BUSCO v6.0.0 and gfastats v1.3.11, the Hi-C-scaffolded Flye assembly was selected as the final assembly based on its superior overall quality and completeness (Supplementary Table S1).

The assembled and annotated genome was subjected to manual curation to further enhance its quality, following the approach routinely used by the Darwin Tree of Life initiative (Howe et al., 2021). Manual inspection enables the detection and correction of misassembled scaffolds as well as the identification and removal of duplicated contigs, thereby improving the structural accuracy of the final chromosomal assembly. This process employed PretextView v1.0.3 (https://github.com/sanger-tol/PretextView/) together with the rapid curation workflow (https://github.com/sanger-tol/curationpretext). Genome quality was assessed after the curation with BUSCO v6.0.0 and gfastats v1.3.11. Furthermore, Kmer-based analysis of the raw reads was performed running jellyfish v2.3.1 (Marçais & Kingsford, 2011) and GenomeScope2.0. (Ranallo-Benavidez, Jaron, & Schatz, 2020), while kmer completeness of the final assembly was assessed using Meryl v.1.4.1 and Merqury v1.3 (Arang Rhie, Walenz, Koren, & Phillippy, 2020). We used a length of k=22 for all k-mer based analyses.

### 2.4. Repeat, gene and telomere annotation

Repeat annotation was carried out generating a custom repeat library with RepeatModeler v2.0.7 (Flynn et al., 2020), then running RepeatMasker v4.2.3 with the *-xsmall* option to soft-mask the genome. Genes were annotated using ANNEVO (Zhang et al., 2026) with default parameters. The annotation completeness was assessed running BUSCO v6.0.0, using the carnivora_odb10 lineage dataset with default parameters. Telomer sequences were annotated using the Telomere Identification ToolKit (tidk v0.2.0) (M. R. Brown, Manuel Gonzalez de La Rosa, & Blaxter, 2025), by first identifying telomeric sequences with tidk explore, then running tidk search.

A six-track circos plot of the nuclear assembly was generated using the *circlize R* package (0.4.18 version) (https://github.com/jokergoo/circlize?tab=readme-ov-file)(Gu, Gu, Eils, Schlesner, & Brors, 2014). The tracks included chromosome ring, telomere positions, gene density, simple repeat content, transposable element content and GC content. Repeat content was calculated for simple repeats and transposable elements as the total number of annotated repeat bases overlapping each 500 kb window, expressed as a fraction of window length. Gene density was estimated as the number of annotated genes overlapping each 500 kb window. GC content was computed for each 500 kb window from the reference assembly sequence. Telomere positions were represented using a three-level confidence scheme, combining interval-based high-confidence telomere calls with chromosome-end telomeric-repeat data.

### 2.5. Mitochondrial genome assembly

The mitochondrial genome was assembled from the raw PacBio HiFi and the genome assembly reads using MitoHiFi v3.2.3 (Allio et al., 2020; Uliano-Silva et al., 2023). Out of the potential assemblies, the most complete and successfully circularized one was selected as the final one. The mitochondrial genome of a Baltic ringed seal (NCBI: NC_008433.1) was used as a reference for the MitoHiFi run.

### 2.6. Synteny analyses

We assessed genome synteny to the available pinniped chromosome-level reference genomes: Baikal seal (*Pusa sibirica*; GenBank: GCA_028975605.1; Yakupova et al., 2023), grey seal (*Halichoerus grypus*; GenBank: GCA_964656455.1), Mediterranean monk seal (*Monachus monachus*; GenBank: GCA_976986665.1), Hawaiian monk seal (*Neomonachus schauinslandiand*; GenBank: GCA_002201575.2), Californian sea lion (*Zalophus californianus*; GenBank: GCF_009762305.2), and the Weddell seal (*Leptonychotes weddellii*; GenBank: GCA_000349705.1) using JupiterPlot v1.1 (https://github.com/JustinChu/JupiterPlot) with the following options: ng =110, m = 10,000,000, MAPQ=10 and linkAlpha = 2. The Synteny and Rearrangement Identified (SyRI) tool was used to further visualize and characterize structural variations (SV) (Goel, Sun, Jiao, & Schneeberger, 2019), through visualization with plotsr (https://github.com/schneebergerlab/syri) (Goel & Schneeberger, 2022). Alignment files for SyRI synteny analysis were created using Minimap2 (Li, 2018). Single chromosome comparisons were made using the inter-chromosomal (itx) plotting method to explore SV in candidate regions between genomes of interest.

### 2.7. Phylogeny

A phylogenetic tree was then generated for all available high-quality (over 80% complete single-copy BUSCO genes) pinniped assemblies from NCBI. To do so, fasta files were obtained and a supermatrix alignment for the BUSCO genes (BUSCO v5.4.7) generated using BUSCO_phylogenomics (https://github.com/jamiemcg/BUSCO_phylogenomics) with default settings. Next, a phylogenetic tree was reconstructed using iqtree v2.3.4 (Minh et al., 2020), utilising the JTT+F+I+R10 substitution model (chosen according to BIC) and ultrafast bootstrapping (1,000) and an SH-like approximate likelihood ratio test (1,000). Lastly, the phylogenetic tree was visualized using TreeViewer 2.2.0 (Bianchini & Sanchez-Baracaldo, 2024).

### 2.8. Gene family expansion/contraction analysis

To investigate candidate gene families putatively associated with expansion/contraction between the Saimaa ringed seal and the grey seal, a pairwise orthogroup copy-number comparison was conducted. Both species were annotated using the same ANNEVO workflow (2.2 version), excluding chromosome Y from the gene prediction. For each annotated gene locus, the longest predicted coding transcript was retained with AGAT (0.8.1 version) (https://github.com/NBISweden/AGAT/tree/master) and translated into protein sequences. Orthogroups were inferred for both species using OrthoFinder (2.5.5 version) (https://github.com/davidemms/OrthoFinder) (Emms & Kelly, 2019). Orthogroups with an absolute copy-number difference of at least 2 were selected as candidate families. Functional annotation of predicted proteins was performed through Diamond (2.1.6 version, e-value threshold 1e-5) (Buchfink, Xie, & Huson, 2015) against UniProtKB/Swiss-Prot database. Sequences without Swiss-Prot hits were subsequently searched against the NCBI non-redundant protein sequences using Diamond in ultra-sensitive mode with an *e*-value threshold of 1e-3 and a maximum of 50 target sequences.

## 3 Results and Discussion

### 3.1 Genome sequencing and assembly

ONT sequencing generated 2.8M reads with a read length N50 of 32.3kb and a total read output of 60.6 Gb. Based on the estimated genome 2.23 Gb size, the sequencing data provided approximately 19.6× coverage. Hi-C sequencing produced 150 Gb, which were used to scaffold the assembly. Of the assembled and manually curated genome 99.76 % was assigned to 17 chromosome-level scaffolds, corresponding to the expected 15 autosomes and the X and Y sex chromosomes (Beklemisheva et al., 2020) (see Supplementary Table S1 for full assembly statistics). The overall assembly statistics are summarised in a snail plot in Figure 2A, with comparison to Earth BioGenome Project assembly benchmarks (A. Rhie et al., 2021) provided in Table 1. The final assembly exhibited a total length of 2.35Gb with a scaffold N50 of 176.8Mb, and a high completeness (96.4 % k-mer completeness) and consensus quality (Merqury QV = 43.8), and accuracy (99.9958 %) (Figure 2, Supplementary Table S1). BUSCO analysis using the mammalia_odb10 reference set (n = 9,226) identified 99.6 % of the expected gene set (single copy = 98.2 %, duplicated = 1.3 %; Figure 2A).

**Figure 2.**
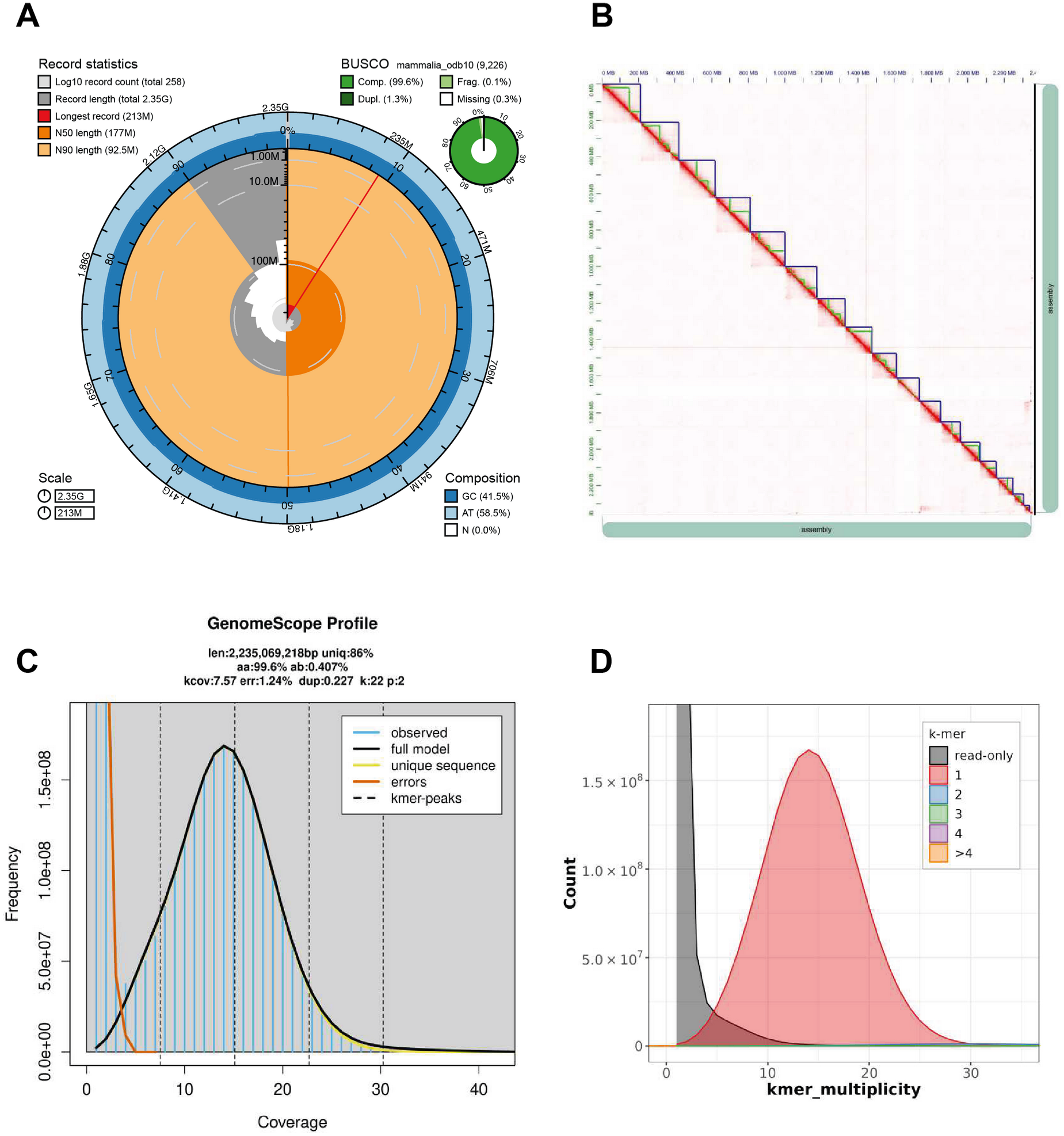
An overview of the Saimaa ringed seal genome assembly mPhs449_Pusa_saimensis_1. (A) The BlobToolKit snail plot summarising the key assembly statistics alongside BUSCO gene completeness. The total circumference corresponds to the full genome length and is divided into 1,000 bins. The outermost blue tracks show the distribution of GC, AT, and N content across these bins. Scaffolds are arranged clockwise in descending order of length and are shown in dark grey, with the longest scaffold highlighted by a red arc. The N50 and N90 scaffold lengths are indicated by darker and lighter orange arcs, respectively. At the centre, a light grey spiral represents the cumulative scaffold count on a logarithmic scale. A summary of complete, duplicated, fragmented, and missing BUSCO genes is displayed in the top-right corner. (B) Hi-C contact map of the Saimaa ringed seal genome assembly. The heatmap visualises chromatin interaction frequencies across the assembled genome, with colour intensity reflecting contact density between genomic regions. Chromosomes (blue boxes) are ordered by decreasing size and labelled along both axes, and a megabase scale is provided on the left and above. Strong interaction signals along the diagonal indicate well-resolved, contiguous chromosome assemblies, while off-diagonal patterns may reflect inter-chromosomal contacts or structural features. (C) Frequency of k-mer distribution generated using GenomeScope2, showing genome size estimate, heterozygosity and repeat content based on unassembled sequencing reads. There is no observable secondary (heterozygous) peak at haploid coverage, underlining the high rate of homozygosity and low genetic diversity of the species. (D) Evaluation of k-mer completeness by MerquryFK, showing the recovery of k-mers from the original read data in the final assemblies. The grey curve is unassembled k-mers in read data and other colors found in other assemblies. Only one haplotype was assembled (red).

**Table 1.**
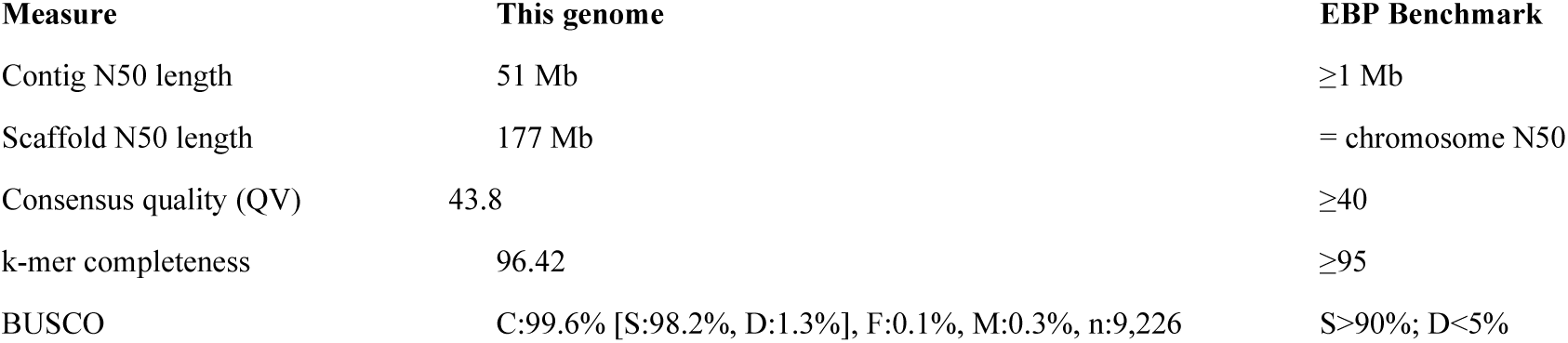
Comparison of the Saimaa ringed seal genome assembly summary metrics with the Earth BioGenome Project (EBP) benchmarks.

The sex chromosomes were identified based on coverage and the Hi-C signal. The two sex chromosomes, originally merged by the assembler at the homologous pseudoautosomal region, were manually separated during curation. The pseudoautosomal region was left as a part of the X-chromosome. Due to the repetitive nature of the Y-chromosome, the rest of its assembly is provided as a single larger scaffold and 25 unlocalised scaffolds (Supplementary Table S1). The final genome assembly is 2.353 Gb long (Supplementary Table S1), is named mPhs449_Pusa_saimensis_1 and is available through NCBI under the accession GCA_059488705.1.

The Saimaa ringed seal genome exhibits features typical of a mammalian genome, including heterogeneous gene density, a substantial proportion of transposable elements, and the presence of simple sequence repeats (Figure 3). Gene-rich regions are generally associated with higher GC content, whereas repeat elements, particularly transposable elements, are enriched in gene-poor regions and contribute significantly to overall genome size and structural complexity. A near telomere-to-telomere assembly was achieved for all chromosomes, with most containing more than 100 copies of the canonical telomeric repeat (TTAGGG) and only Chr2, Chr5, Chr10, as well as ChrY ends remaining less completely resolved. As expected for mammalian genomes, repeat-rich regions such as centromeres and telomeres remain challenging to resolve completely, which may explain residual assembly uncertainties and unplaced scaffolds. Overall, mPhs449_Pusa_saimensis_1 captures the characteristic organisation and composition of a high-quality mammalian genome assembly.

**Figure 3.**
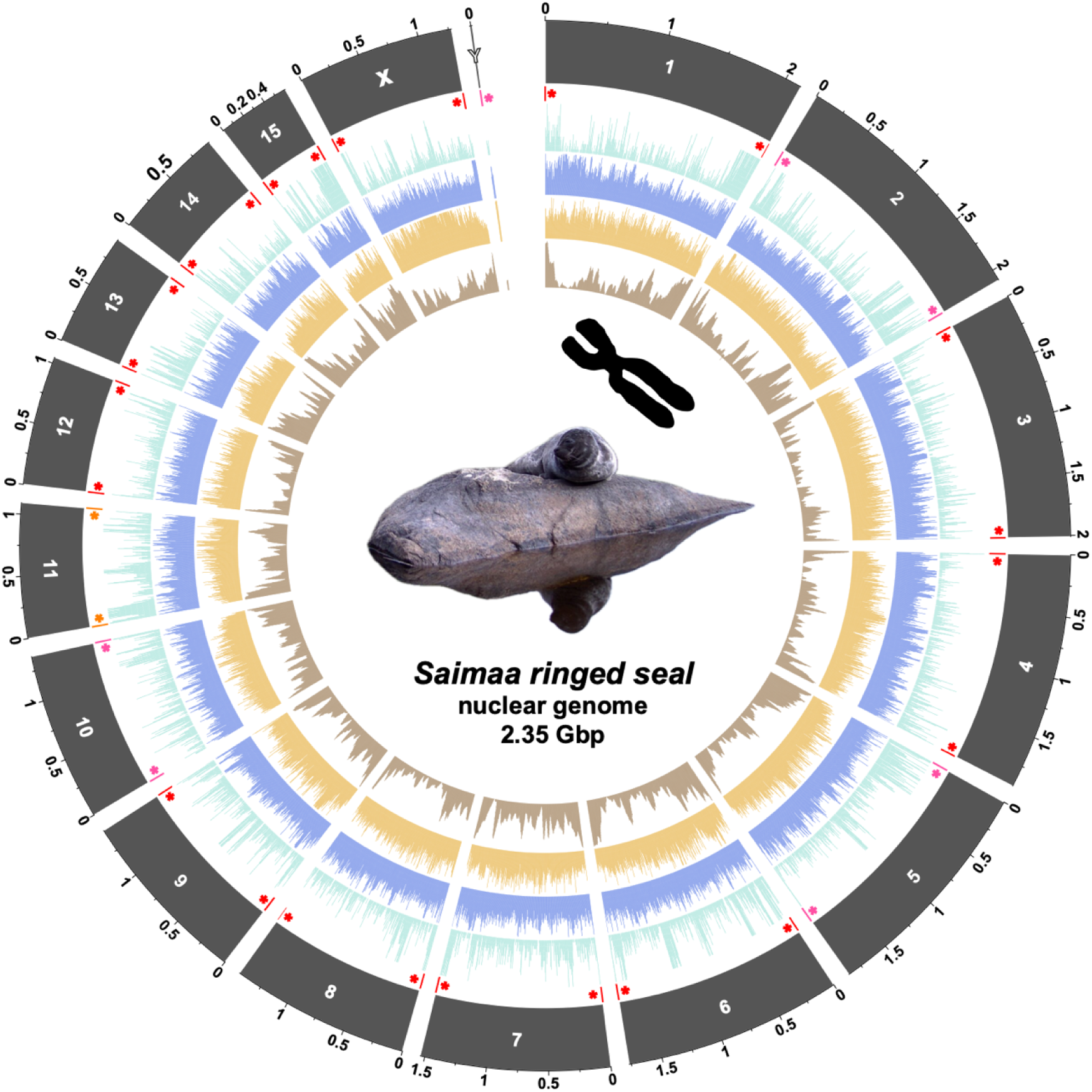
Genome organisation of the Saimaa ringed seal nuclear assembly (mPhs449_Pusa_saimensis_1). The outermost ring shows chromosomes (scale 100 Mb), followed by a track showing telomere positions. The next track (turquoise) represents gene density calculated in 500 kb windows, followed by simple repeat content (blue) and transposable element content (orange). The innermost brown track depicts GC content. To enhance visual contrast, track values were scaled using upper quantile caps. For repeat annotations, the total number of bases overlapping each 500 kb window was calculated separately for simple repeats and transposable elements and expressed as a fraction of the window length. Telomere positions are shown using a three-level confidence scheme: high-confidence calls based on interval detection with repeat counts greater than 100 (red asterisks), medium-confidence support from chromosome-end windows exceeding a repeat threshold of 20 (orange asterisks), and low-confidence support from chromosome-end windows exceeding a threshold of 5 (pink asterisks).

### 3.2 Saimaa ringed seal mitochondrial genome

The mitochondrial genome of the Saimaa ringed seal is 16,821 bp in length and exhibits the canonical gene content and organization characteristic of mammalian mitogenomes (Figure 4A), including 13 protein-coding genes, 22 tRNAs, and two rRNAs arranged in the conserved vertebrate order. No structural rearrangements relative to other phocid seals were detected. In addition to the coding region, we provide a detailed functional annotation of the 1,385 bp non-coding region (NCR) (Figure 4B). Annotation was guided by sequence similarity and structural features conserved in human and mouse mitochondrial DNAs, allowing identification of key regulatory elements within the NCR, including conserved sequence blocks (CSBs) and termination-associated sequences (TAS). Haplotype-differentiating SNPs are located primarily within hypervariable segments I and II (HVS-I and HVS-II) at the 3′ end of the displacement loop (D-loop) structure. The mitogenome of the Psh449 corresponds to the most common Saimaa ringed seal haplotype (H3) (Heino et al., 2023).

**Figure 4.**
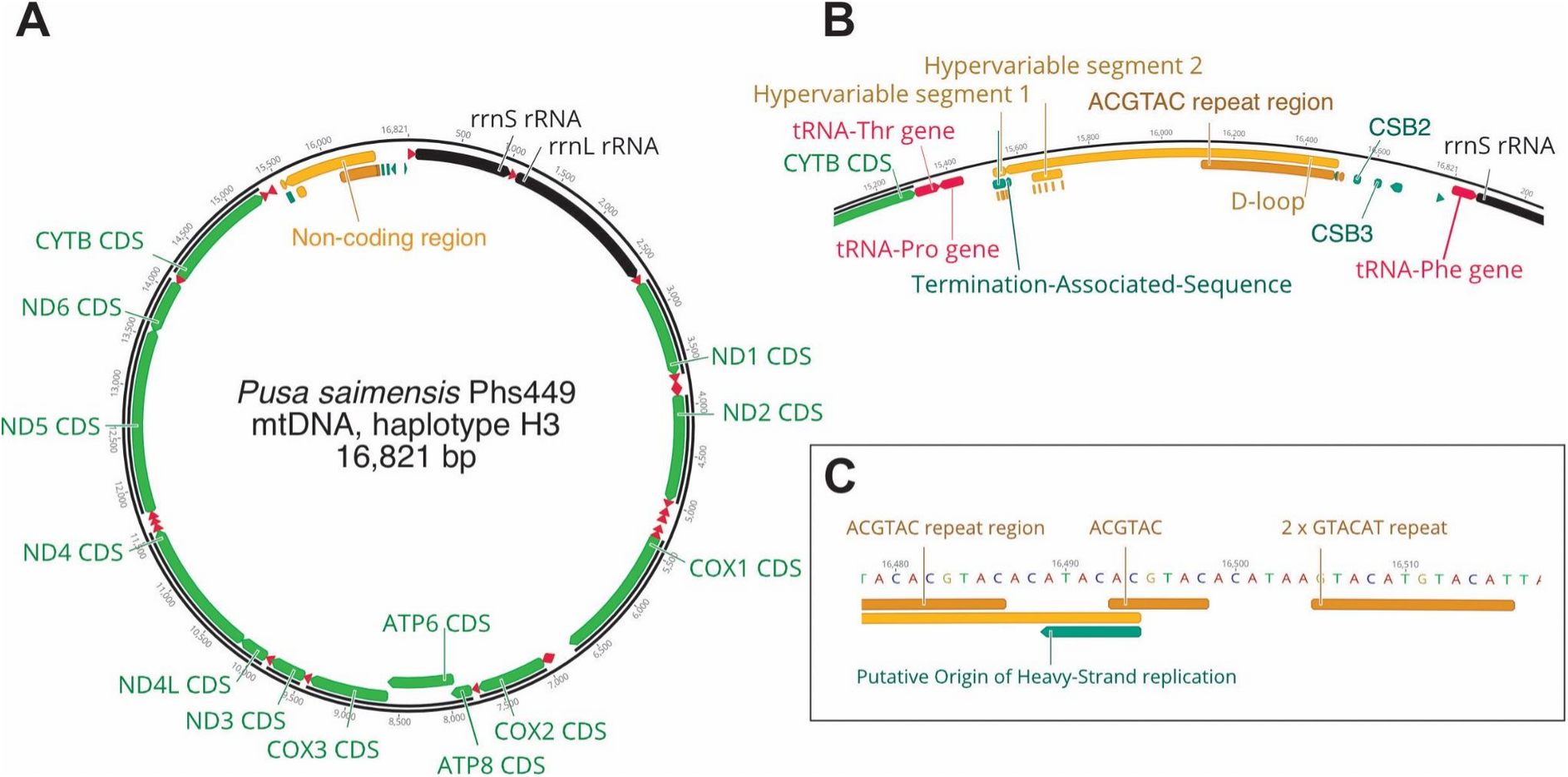
Mitochondrial genome of the Saimaa ringed seal Phs449. (A) Circular representation of the mitochondrial genome. Protein-coding genes are shown in green, tRNA genes in red and rRNA genes in black. The 1,385 bp non-coding region (NCR) is highlighted. (B) Examples of the functional annotations within the NCR. Besides putative control elements, the region contains an array of 63 tandem repeats of the hexanucleotide motif ACGTA, as well as (C) an additional isolated stretch of the same hexamer and its partial iteration.

The NCR of Psh449 contains a 378 bp short repeat (SR) region composed predominantly of 63 tandem copies of the hexanucleotide motif ACGTAC, together with minor sequence variants, including two GTACAT repeat units separated by short intervening nucleotides (Figure 4C). Similar SR arrays have been described in other phocid (Arnason, Gullberg, Johnsson, & Ledje, 1993; Arnason & Johnsson, 1992) and otariid seals (Hoelzel, Hancock, & Dover, 1993) and appear to represent a common feature of carnivoran mitogenomes (Hoelzel, Lopez, Dover, & O’Brien, 1994). Similar short tandem repeats are also present in lagomorphs such as hares, which also have longer tandem repeat arrays of ∼190 bp units (Tapanainen et al., 2024).

The functional significance of these short repeat regions remains unclear. However, their repetitive structure makes them prone to replication slippage, leading to frequent length variation through expansion and contraction (Pfeuty, Guéride, & Lecellier, 2001). Mutation rates in such regions have been estimated to reach up to 10^-2^ per generation (Casane, Dennebouy, deRochambeau, Mounolou, & Monnerot, 1997). Given the exceptionally low haplotypic diversity of the Saimaa ringed seal population (Heino et al., 2023; Olsen et al., 2025), length polymorphism within the SR array may provide a useful supplementary marker for population genotyping and monitoring.

The mitochondrial DNA (mtDNA) sequence difference between the Saimaa ringed seal and the Baltic ringed seal (*Pusa hispida*) was relatively high (2.12%). Using the “universal” molecular clock for mtDNA (2% sequence divergence per million years)(W. M. Brown, George, & Wilson, 1979), this would place the divergence of the two lineages in the mid-Pleistocene 1Ma ago. However, we would not put too much emphasis on this timing, as substantial variation in mitochondrial substitution rates is well documented across taxa and even among closely related lineages (Nabholz, Glemin, & Galtier, 2008, 2009). It is noteworthy that mitochondrial genomes of ringed seals exhibit high standing diversity (Olsen et al., 2025), which is likely influenced by historically large effective population sizes that allow the persistence of deep coalescent lineages (Löytynoja et al., 2025). In addition, life-history traits such as metabolic rate can affect mutation rates in mtDNA, potentially leading to rate heterogeneity among lineages (Nabholz et al., 2008, 2009). Differences in generation time, demographic history (e.g. bottlenecks and expansions), and the action of purifying selection on mitochondrial genes can further distort simple clock-based estimates. Nevertheless, the observed level of mitochondrial divergence is consistent with the previously inferred deep evolutionary origins of the Saimaa ringed seal lineage.

### 3.3. Chromosomal architecture of the Saimaa ringed seal

The presented assembly showed only minor synteny differences in its chromosomal architecture compared to other available pinniped chromosome-level genome assemblies, even with an otariid species (Figure 5). The analysis also confirmed the previously reported evolutionarily highly conserved genome structure among pinnipeds (Beklemisheva et al., 2020). In particular, the cytogenetically described apomorphic fusion events in Phocini and Phocidae (Beklemisheva et al., 2020) are consistent with our sequence-level results, corresponding to Chromosomes 2 and 7, respectively (Figure 5). Another noteworthy interchromosomal rearrangement includes a translocation between Chromosomes 6 and 7, when comparing phocid species with the otariid Californian sea lion. While showing generally a low number of interchromosomal rearrangements among pinnipeds, more distant taxa, such as both extant monk seal species and the sea lion, show an increasing degree of intrachromosomal rearrangements compared to the Saimaa seal (Figure 5). A large-scale intrachromosomal rearrangement affecting chromosome 2 seems to be apomorphic for the Saimaa ringed seal. However, this region warrants cautious interpretation. During manual curation using PretextView (see Methods), a segment of chromosome 2 exhibited notably weak Hi-C contact signals with the remainder of the chromosome, and its orientation and placement could not be unambiguously resolved despite extensive inspection. Although no clear assembly conflicts or misplaced telomeric signals were detected, the possibility remains that this segment may be misassembled or incorrectly oriented (Figure 5 & 6A,B). It could thus either represent a genuine lineage-specific rearrangement or a structurally labile region that is difficult to assemble.

**Figure 5:**
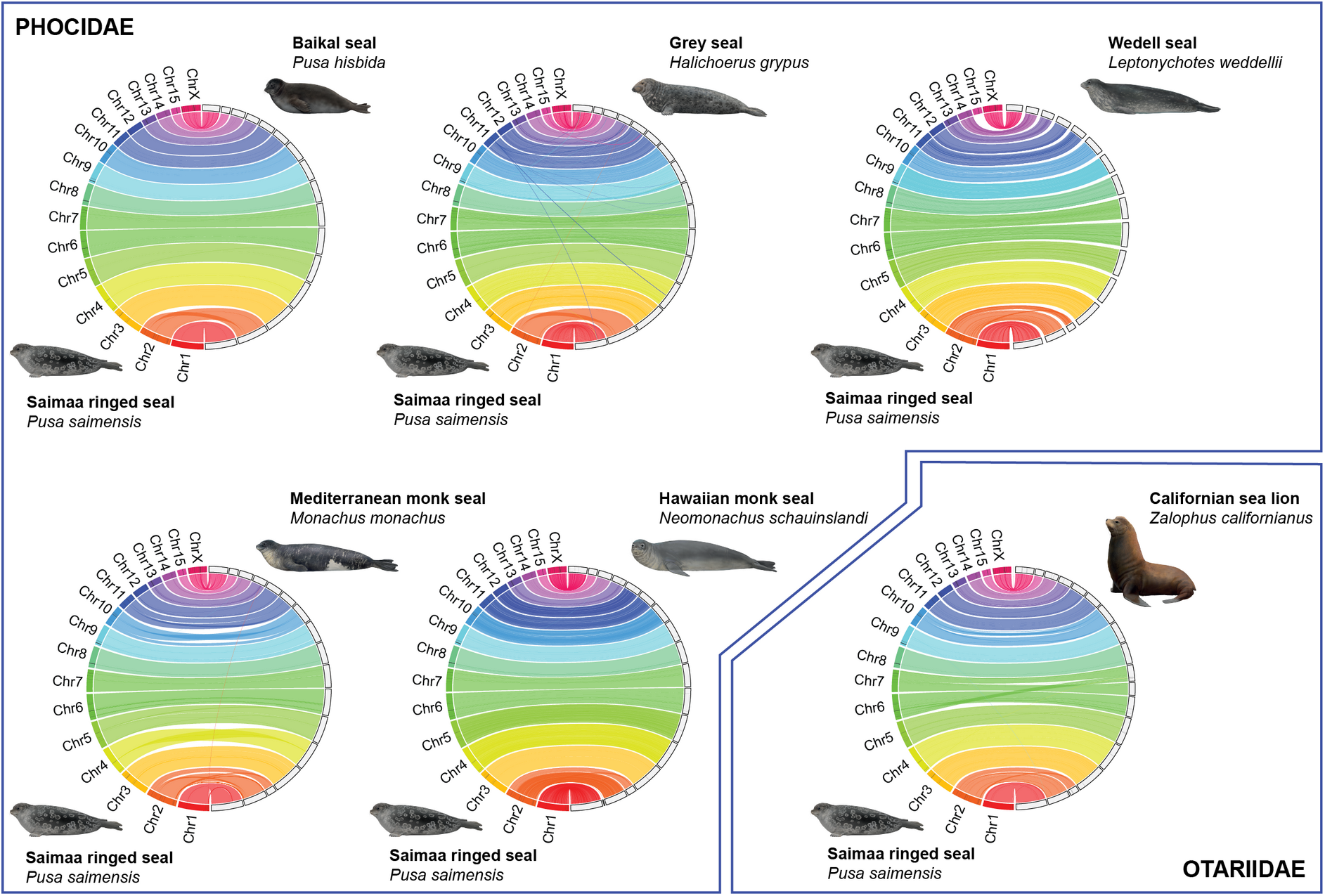
Circos plots comparing the synteny of the chromosome-scale genome assembly of Pusa saimensis (mPhs449_Pusa_saimensis_1) with chromosome-scale assemblies of other pinniped species. These include another freshwater species, the Baikal seal (Pusa sibirica); two European phocids, the grey seal (Halichoerus grypus) and the Mediterranean monk seal (Monachus monachus); and one Antarctic marine phocid, the Weddell seal (Leptonychotes weddellii). The Californian sea lion (Zalophus californianus) represents an otariid species. A notable structural rearrangement is apparent in chromosome 2 of P. saimensis, and chromosome 15 is absent from some of the assemblies.

We next sought to investigate chromosome-level differences in more detail using SyRI. In several pairwise comparisons, some chromosomes were excluded from downstream visualization because no usable compatible syntenic or structural variation records remained after filtering. As these chromosomes are not absent in the assemblies, the results are likely to reflect cases where (i) no syntenic regions were retained by SyRI, (ii) detected variants did not belong to the classes included in the plotting step, or (iii) alignments were too fragmented or weak to pass filtering to the applied filtering criteria. Consequently, some chromosomes are present in the assemblies but not represented in the final plotting files. The extent of missing chromosome comparisons varied markedly among pinniped species. The most extreme case was the Weddell seal, for which multiple chromosomes (Chr2–7 and Chr15) were not retained, followed by the Mediterranean monk seal, the Hawaiian monk seal and the Californian sea lion, each lacking several chromosomes in pairwise comparisons. In contrast, the Baikal seal showed missing data for only Chr1 and Chr2 and the grey seal for Chr8 and Chr9. Given these limitations, we focused subsequent analyses on the grey seal, which is both evolutionarily close to the Saimaa ringed seal and yielded the most complete and reliable pairwise comparison. Importantly, the two chromosomes missing in this comparison are relatively small, especially when contrasted with Chr1 and Chr2, which are absent in the Baikal seal comparison. Furthermore, the analyses also showed the large rearrangement previously observed in the Jupiter plots (Figure 6A and B).

**Figure 6:**
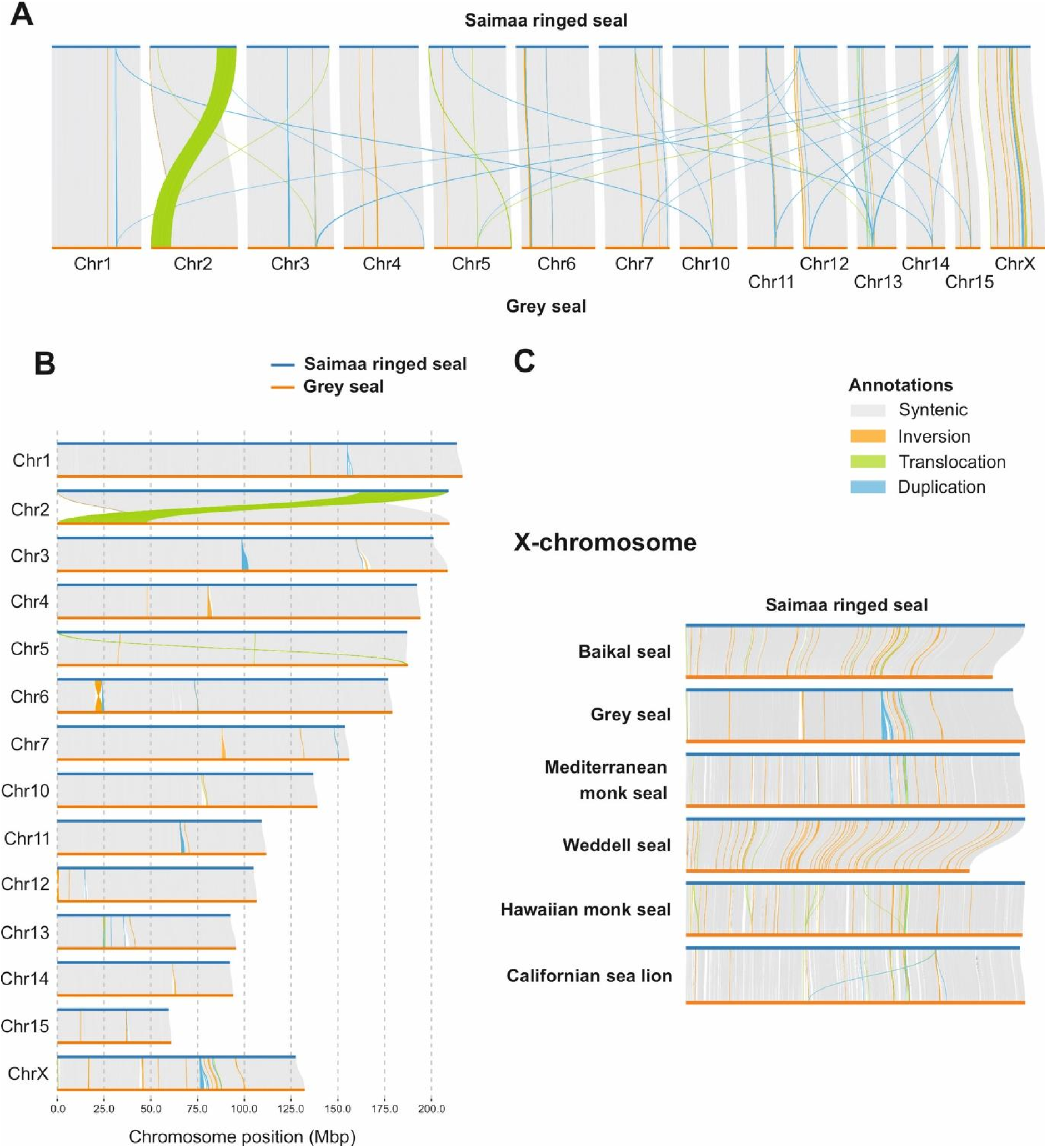
SyRI-based synteny analysis of the Saimaa ringed seal genome assembly. (A) Pairwise comparison between Saimaa ringed seal and grey seal chromosomes, highlighting inter-and intrachromosomal rearrangements. Note the large translocation between Chr2 ends. (B) Detailed view of intrachromosomal differences between Saimaa ringed seal and grey seal, including a short translocation at the ends of Chr5 and a larger inversion on Chr6. (C) Comparison of ChrX across pinniped reference genome assemblies, showing that most differences are attributable to inversions.

For the X chromosome, sufficient synteny was conserved to enable cross-species analysis (Figure 6C). Structural rearrangements and other sequence variants identified by SyRI for the X chromosome are summarized in Table 2. It should be noted that X-chromosome evolution is faster than for the autosomes, due to the phenomenon called “the faster-X effect” (Meisel & Connallon, 2013). Besides true biological variation, this analysis also reflects differences between reference genome assemblies.

**Table 2.** Comparison of the Saimaa ringed seal X chromosome assembly with other pinniped reference genome assemblies. DUP: duplications; INV: inversions; INVDP: inverted duplications; INVTR: inverted translocations; SYN: total length of syntenic regions; TRANS: translocations; DEL: deletions; INS: insertions; SNP: single-nucleotide polymorphisms. Higher SYN values indicate lower collinearity (synteny) between chromosome assemblies.

| Species | DUP | INV | INVDP | INVTR | SYN | TRANS | DEL | INS | SNP |
| --- | --- | --- | --- | --- | --- | --- | --- | --- | --- |
| Baikal seal | 1 | 41 | 0 | 8 | 53 | 4 | 40,183 | 48,900 | 603,682 |
| Grey seal | 55 | 28 | 70 | 5 | 36 | 5 | 51,865 | 51,794 | 381,045 |
| Weddell seal | 1 | 86 | 1 | 2 | 87 | 2 | 111,284 | 104,204 | 1,415,644 |
| Mediterranean monk seal | 1 | 66 | 9 | 8 | 78 | 2 | 105,153 | 102,585 | 1,357,869 |
| Hawaiian monk seal | 0 | 79 | 6 | 16 | 103 | 9 | 107,347 | 101,035 | 1,379,529 |
| California sea lion | 11 | 67 | 14 | 16 | 91 | 12 | 94,039 | 94,590 | 1,441,575 |

During assembly the X and Y chromosomes were originally merged by the assembler at the pseudoautosomal region and subsequently separated during manual curation. At this step, we decided to not duplicate this region, as it would only cause reads to randomly split between the two chromosomes during an alignment. Thus, the pseudoautosomal region is only included as part of the X chromosome. This part consistently showed diploid-like coverage in resequencing and alignment of male individuals and includes a telomeric region at one end.

Interestingly, the grey seal shows fewer SNP differences relative to the Saimaa ringed seal than does the congeneric Baikal seal. At face value, this could suggest a closer evolutionary affinity between the grey seal and Saimaa ringed seal. However, the phyletic relationships among phocidae are notoriously complicated (Higdon, Bininda-Emonds, Beck, & Ferguson, 2007). Although some analyses place *Halichoerus* within the *Pusa* clade, this is not robust but appears to be driven by compositional or rate heterogeneity and limited sampling (Fulton & Strobeck, 2010; Higdon et al., 2007; Nebenführ, Arnason, & Janke, 2024).

### 3.4. Phylogenomics of pinnipeds based on single-copy orthologues

The genome-wide phylogenetic analysis recovered a well-resolved and fully supported topology across 17 pinniped species (Figure 7). The placement of the *Pusa saimensis* was consistent with expectations, forming a sister relationship with *Pusa hispida botnica*. Despite this close relationship, the two lineages were clearly separated, with maximal support (ultrafast bootstrap = 100), reflecting their distinct evolutionary trajectories (Löytynoja et al., 2025). The major pinniped families and subfamilies were recovered as monophyletic and highly congruent with previously published phylogenies (Fulton & Strobeck, 2010; Higdon et al., 2007; Nebenführ et al., 2024; Zhao, Yu, Yang, Seim, & Tian, 2026). Consistent with the recent genome-scale phylogeny (Zhao et al., 2026), *Halichoerus grypus* was recovered as the sister lineage to the *Pusa* clade, while *Phoca vitulina* formed their sister lineage. The agreement across these independent phylogenetic approaches demonstrates that conserved single-copy BUSCO genes provide robust phylogenomic inference for resolving relationships across Pinnipedia. Species with higher levels of missing data, such as the *Leptonychotes weddellii* and the *Pusa sibirica* (see BUSCO completeness values in Figure 7), exhibited comparatively longer branch lengths. This pattern is consistent with reduced gene recovery inflating apparent divergence in concatenation-based analyses (Xi, Liu, & Davis, 2016) and highlights the importance of genome completeness in phylogenomic inference. Nevertheless, these taxa retained their expected phylogenetic positions with maximal support, indicating that the overall topology is robust despite differences in assembly completeness.

**Figure 7.**
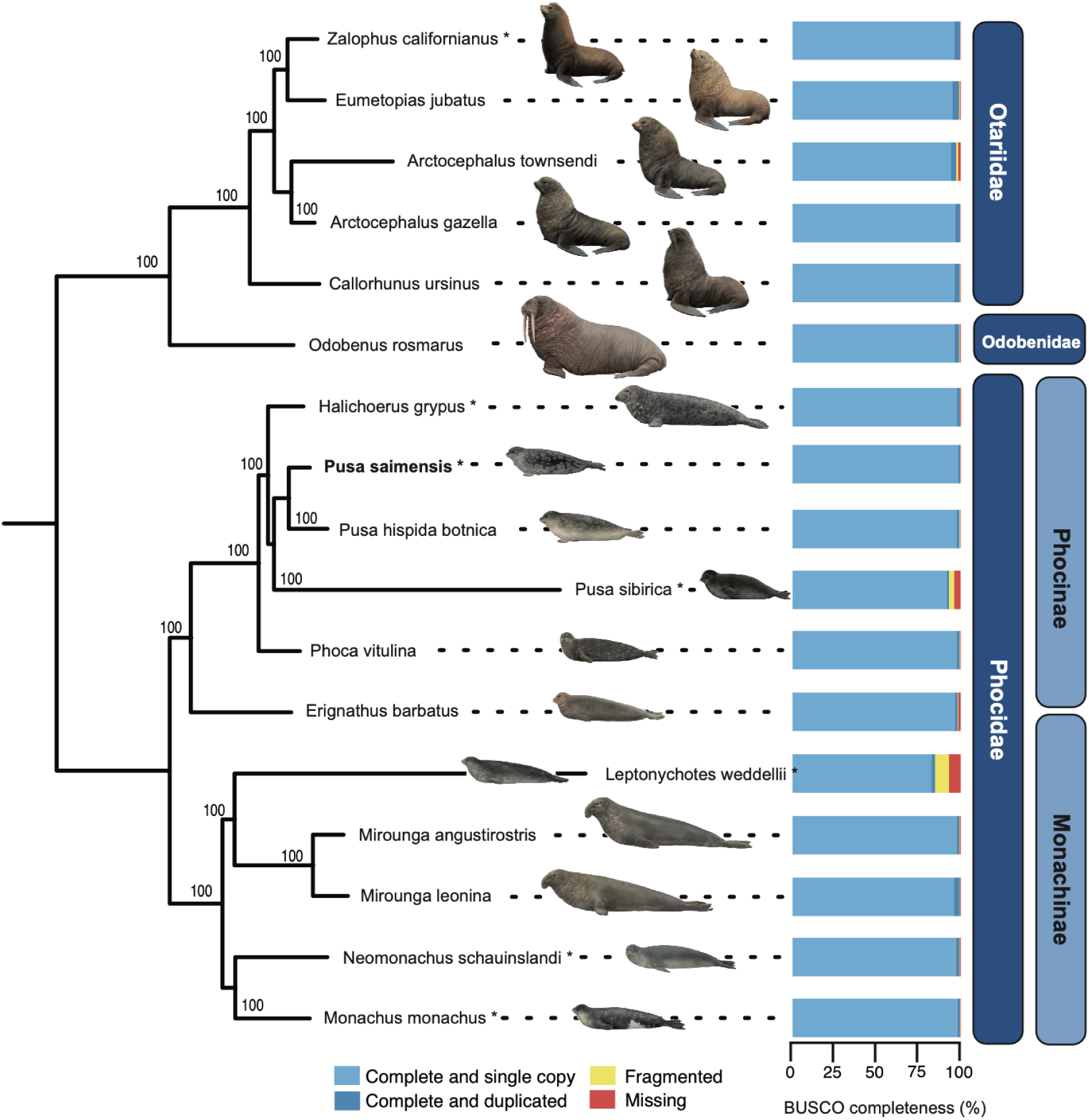
Coalescent phylogenetic tree from 17 pinniped species based on 9,226 BUSCO genes, each with the assigned BUSCO completeness [%]. Saimaa ringed seal is marked in bold and other species with available chromosome level genome assemblies with an asterisk (*). Family-and subfamily-level taxon assignments are indicated. Seal images © Lynx Nature Books, illustrated by Toni Liobet.

### 3.5. Lineage-specific gene family expansions and contractions in Saimaa ringed seal

To identify genomic features potentially associated with lineage-specific adaptation, we compared gene family expansions and contractions between the Saimaa ringed seal and its closest marine relative with a reference-quality genome assembly (Figure 7), the grey seal. For both species, the longest predicted coding transcript per locus was retained and translated into protein sequences, yielding high-quality and highly representative protein sets. Pairwise orthogroup analysis identified a total of 17,799 orthogroups. The majority of these gene families were conserved in copy number between the two species, with 17,043 orthogroups showing identical counts. In contrast, 91 orthogroups showed higher copy number in *P. saimensis*, whereas 97 showed higher copy number in *H. grypus* (Supplementary Table S2). These results indicate that most gene families are stable between these species, although a limited subset displays asymmetry in copy number.

Functional annotation of genes within the differing orthogroups revealed both shared and species-specific gene classes (Supplementary Table S2). Closer inspection showed that many of these orthogroups contained homologous gene families that had likely been split into separate groups due to sequence divergence between the species. After consolidating such cases, additional orthogroups with effectively identical copy numbers could be excluded. The remaining gene families were grouped into 12 functional categories (Figure 8).

**Figure 8.**
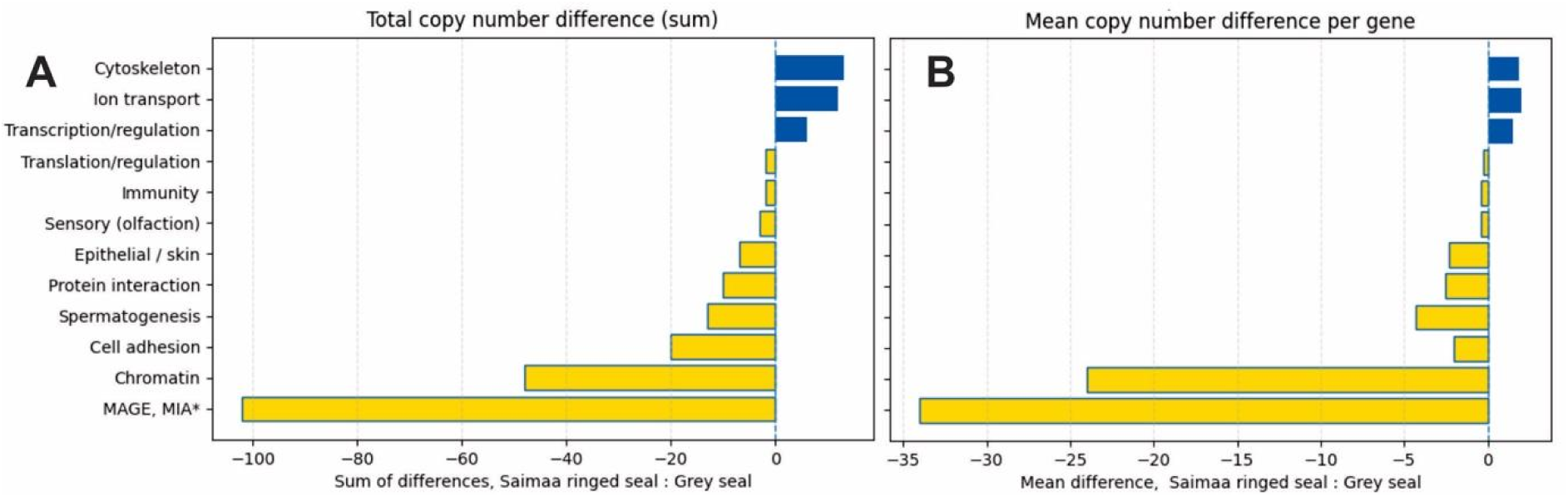
Pairwise comparison of the Saimaa ringed seal and the grey seal, illustrating lineage-specific expansion or contraction of gene families. (A) Sum and (B) mean copy number differences per functional category. * Melanoma-associated antigen (MAGE) and melanoma inhibitory activity (MIA) gene families.

Among these categories, only cytoskeleton-, ion transport-, and transcription/regulation-related genes showed higher copy numbers in the Saimaa ringed seal compared to grey seal (Figure 8). The ion transport category consisted primarily of voltage-gated sodium channel genes. Although the functional significance of this expansion is unclear, it might reflect broader modifications of ion homeostasis associated with adaptation to freshwater conditions. Freshwater mammals experience continuous osmotic influx of water and passive ion loss, requiring efficient conservation of sodium and other electrolytes (Evans, 2023). Previous comparative genomic analyses of the Saimaa ringed seal and the Baikal seal have identified adaptive changes in multiple genes and regulatory elements involved in renal development, the renin-angiotensin system, sodium transport and epithelial ion handling (Zhao et al., 2026), suggesting that freshwater adaptation involves coordinated evolution of osmoregulatory pathways rather than changes in individual transport proteins alone. In this context, the expansion of sodium channel genes reported here may represent one component of a broader genomic response supporting ion homeostasis. It is noteworthy that the Saimaa ringed seal kidneys also differ in their morphology from those of the marine seals (Nihtilä & Laakkonen, 2025). Similarly, variation in genes related to cytoskeleton, cell adhesion, and epithelial structure may reflect differences in mechanical or physiological demands between freshwater and marine environments (Zhao et al., 2026).

In contrast, the grey seal showed higher copy numbers in several other categories. Remarkably, a pronounced expansion of melanoma-associated antigen (MAGE) and melanoma inhibitory activity (MIA) gene families. Although these genes are primarily studied in the context of tumors in humans, they are part of rapidly evolving, often immune-or spermatogenesis-related mammalian gene families and may play broader roles in host-pathogen interactions, reproduction or cellular stress responses (Katsura & Satta, 2011; Tacer et al., 2019). The grey seal also showed higher copy numbers of certain spermatogenesis-related genes. This is notable given differences in mating systems: grey seals exhibit a polygynous breeding system, in which dominant males monopolise access to multiple females (Anderson & Fedak, 1985; Twiss, Cairns, Culloch, Richards, & Pomeroy, 2012), whereas ringed seals are more spatially dispersed and exhibit less pronounced sexual competition (Le Boeuf, 1991; Stirling, 1983). Expansion of spermatogenesis-related genes, including the MAGE/MIA-genes described above, in grey seals may therefore be linked to increased reproductive competition. The reason for more numerous histone H2B variants in grey seal is less obvious. Histone variants usually replace canonical histones for specific tasks of genome maintenance and heterochromatin formation in different tissues or developmental stages (Talbert & Henikoff, 2017). It is tempting to speculate that their expansion could correlate with the MAGE/MIA functions in grey seals. Furthermore, olfactory receptor (OR) genes were reduced in the Saimaa ringed seal compared to the grey seal. OR gene families are known to undergo rapid birth and death evolution, with gene duplication and loss closely tracking ecological adaptation (Niimura, 2012). In marine mammals, reduced reliance on olfaction has been associated with relaxed selection and extensive OR gene loss (Liu et al., 2019). The additional reduction observed in Saimaa ringed seal may therefore represent a lineage-specific consequence of adaptation to the ecologically simplified freshwater environment of Lake Saimaa, where a more limited prey spectrum may have reduced the functional constraints of OR diversity maintenance.

Taken together, these results indicate that gene copy number differences between the Saimaa ringed seal and grey seal are primarily driven by a limited number of gene families, including rapidly evolving immune-and reproduction-related genes, as well as sensory and structural components. While these patterns remain correlative, they provide biologically plausible hypotheses for future comparative studies on the physiological and ecological adaptations of freshwater and marine pinnipeds.

## 4 Conclusion

The presented chromosome-level reference genome for the Saimaa ringed seal substantially advances genomic resources for pinnipeds and for this highly endangered endemic freshwater seal. It enabled a first view into potential mechanisms of its evolution and genomic architectural changes in pinnipeds, which can establish a foundation for future functional and comparative studies. Being an endangered species, this reference genome enables high-resolution investigations into its genetic diversity, inbreeding, and demographic history, providing critical tools for its conservation and long-term management.

## Author Contributions

**Martin Grethlein:** data curation (equal), formal analysis (equal), investigation (equal), methodology (equal), writing – original draft (equal), writing – review and editing (equal). **Zsófia Fekete:** data curation (equal), formal analysis (equal), investigation (equal), methodology (equal), validation (equal), visualization (equal), writing – original draft (equal), writing – review and editing (equal). **Steffi Goffart:** investigation (equal), resources (equal), writing – original draft (equal), writing – review and editing (equal). **Angelika Kiebler:** formal analysis (equal), writing – original draft (equal), writing – review and editing (equal). **Mervi Kunnasranta:** conceptualization (equal), writing – review and editing (equal). **Marja Niemi:** investigation (equal), writing – review and editing (equal). **Danilo F. Santoro:** conceptualization (equal), data curation (equal), formal analysis (equal), investigation (equal), methodology (equal), visualization (equal), writing – original draft (equal), writing – review and editing (equal). **Gerrit Wehrenberg:** formal analysis (equal), investigation (equal), methodology (equal), validation (equal), visualization (equal), writing – original draft (equal), writing – review and editing (equal). **Sven Winter:** formal analysis (equal), writing – review and editing (equal). **Stefan Prost:** conceptualization (lead), data curation (equal), formal analysis (equal), funding acquisition (equal), methodology (equal), project administration (equal), supervision (lead), visualization (equal), writing – original draft (equal), writing – review and editing (lead). **Jaakko Pohjoismäki:** conceptualization (lead), data curation (equal), formal analysis (equal), funding acquisition (equal), methodology (equal), project administration (lead), supervision (equal), visualization (equal), writing – original draft (lead), writing – review and editing (lead).

## Supporting information

**Supplementary Table S1**: Genome assembly statistics and quality metrics.

**Supplementary Table S2**: Orthogroup assignments and differences between *P. saimensis* and *H. grypus*.

## Acknowledgements

We thank Miina Auttila, Riikka Alakoski and Mikko Suonio from Metsähallitus Parks & Wildlife Finland for their essential fieldwork and assistance in sample recovery. We are grateful to veterinarian Sanna Sainmaa (Korkeasaari Zoo, Helsinki, Finland) for performing the biopsy procedure that enabled us to obtain the fibroblast cell line. Dr Dominic Absolon (Wellcome Sanger Institute, Tree of Life Programme, Hinxton, UK) is thanked for providing quality control for the manual curation of the assembly. We are grateful to Dr Camila Di Nizo and Dr Jonas Astrin (ZFMK, Bonn, Germany) for the biobanking of the cell line. Ari Löytynoja and Emmi Olkkonen (University of Helsinki), as well as Tommi Nyman (NIBIO, Svanhovd, Norway) are thanked for the useful discussions on Saimaa ringed seal genomics. Research has received funding from the European Union’s LIFE programme and projects Our Saimaa Seal LIFE (LIFE19 NAT/FI/000832) and Support of Survival LIFE (101292872 LIFE25-NAT-FI-LIFE SOS). The material reflects the views of the authors, neither the European Commission nor the CINEA is responsible for any use that may be made of the information it contains. Martin Grethlein is funded by the Onni Talas Foundation. Stefan Prost and Gerrit Wehrenberg are funded by the University of Oulu and the Research Council of Finland Profi6 336449 program “Biodiverse Anthropocenes.”

## Data Availability Statement

The genome assembly has been deposited in NCBI GenBank under accession number GCA_059488705.1. The associated project and sample metadata are available under BioProject accession PRJNA1450854 and BioSample accession SAMN57173219, respectively.

