## Supplementary Table S1 for "Chromosome-level reference genome assembly of the Saimaa ringed seal (*Pusa saimensis*) – an ancient glacial relict landlocked pinniped"

| **BUSCO** | **Flye + HiC** | **Hifiasm + HiC** | **Final Curated** |
| --- | --- | --- | --- |
| Complete | 9,186 (99.6%) | 9,028 (97.9%) | 9,186 (99.6%) |
| Complete and single-copy | 9,061 (98.2%) | 8,510 (92.2%) | 9,062 (98.2%) |
| Complete and duplicated | 125 (1.4%) | 518 (5.6%) | 124 (1.3%) |
| Fragmented | 10 (0.1%) | 34 (0.4%) | 10 (0.1%) |
| Missing | 30 (0.3%) | 164 (1.8%) | 30 (0.3%) |
| Total | 9226 | 9226 | 9226 |
| **ASSEMBLY STATISTICS** |  |  |  |
| Number of contigs/scaffolds | 362 | 1,084 | 252 |
| L50 | 7 | 7 | 6 |
| N50 | 153.4e6 | 155.5e6 | 176.8e6 |
| Max Length Contig | 213.4e6 | 205.7e6 | 213.4e6 |
| Total Length | 2.353e9 | 2.527e9 | 2.353e9 |
